# Superiority Illusion in Older Adults: Volume and Functional Connectivity of the Precuneus

**DOI:** 10.1101/2024.09.13.612998

**Authors:** Yuki Shidei, Daisuke Matsuyoshi, Ayako Isato, Genichi Sugihara, Hidehiko Takahashi, Makiko Yamada

## Abstract

**Aim:** Positive thinking, which has been known to extend the lifespan, tends to increase with age. One specific form of positive thinking, the superiority illusion (SI), involves the belief that one is better than others. Despite its potential benefits in aging, the neural basis of the SI in elderly populations remains underexplored.

**Methods:** This study combined a behavioral task, voxel-based morphometry (VBM), and resting-state functional connectivity (rsFC) analyses to investigate the neural substrates of the SI in a cohort of 100 participants, including young (N = 33), middle-aged (N = 33), and older adults (N = 34).

**Results:** Our findings indicated that higher SI scores in older adults were correlated with greater gray matter volume in the right precuneus and stronger rsFC between the right precuneus and the left lateral occipital cortex. However, these correlations were not evident in younger and middle-aged groups.

**Conclusions:** Our findings underscore the importance of the right precuneus and its connectivity in the manifestation of the SI, particularly in older adults, highlighting its potential role in adaptive aging processes.

## Introduction

Enhanced mental health has been significantly associated with increased life expectancy, a correlation well supported by epidemiological studies (1). Indeed, studies have shown that psychological well-being promotes engagement in physical activities and adherence to nutritious diets, thereby augmenting lifespan (2). Conversely, adverse psychological states, such as pessimism and chronic stress, detrimentally influence physiological processes and negatively impact lifestyle choices (3). Positive thinking, a crucial facet of mental health, has been shown to enhance relationship quality and longevity (4). Meanwhile, recent empirical evidence suggests that optimistic individuals likely experience a lifespan extension of approximately 5.4% (5).

Healthy older individuals often exhibit a positive bias, which is a psychological propensity to perceive and interpret information favorably (6). This bias is crucial for maintaining psychological well-being, especially as individuals age. The superiority illusion (SI) is particularly noteworthy among the various facets of positive bias. The SI encompasses the belief in one’s superiority over others, such as being better than the average peer in various domains like intelligence, social skills, and personality (7). As such, understanding the neural correlates of the SI and its manifestation in healthy aging can shed light on the adaptive functions of positive thinking in preserving mental health and cognitive resilience later in life.

While research has begun to uncover the neural underpinnings of positive thinking among younger populations, studies focusing on older adults remain scarce. Previous research has indicated a significant positive association between the propensity for optimism and the volume of the anterior cingulate cortex (ACC) (6). Additionally, older adults who perceive themselves as younger exhibit preserved inferior frontal gyrus and superior temporal gyrus volumes (8). In younger adults, a positive correlation between dispositional optimism and the putamen volume has been observed (9). However, the neural basis of positive thinking in relation to aging remains poorly understood.

The present study aimed to elucidate the relationship between the SI and regional brain volume and functional connectivity among adults of different age groups. We employed voxel-based morphometry (VBM) to assess correlations between regional brain volume and the SI, and utilized resting-state functional magnetic resonance imaging (fMRI) to examine functional connectivity. By integrating these approaches, we seek to clarify the relationship between positive mental states and the neuroanatomical and functional properties of the aging brain.

## Methods

### Participants

For this study, 145 healthy volunteers were recruited from the National Institutes for Quantum Science and Technology and subsequently categorized into three age groups: young (<40 years; mean ± SD = 23.8 ± 3.7 years; N = 84, 11 females), middle-aged (40–59 years; 48.9 ± 5.8 years; N = 37, 24 females), and older group (≥60 years; 68.0 ± 4.7 years; N = 24, 10 females). The exclusion criteria included contraindications to MRI, diagnosed neuropsychiatric disorders, a history of significant head trauma, claustrophobia, and metal implants. Before the study, all participants received a full explanation regarding the objectives and methodology of the study, after which informed consent was obtained from them.

### Assessment of SI

The measurement procedure for the SI has been detailed previously (7, 10). Briefly, 52 traits identified in the literature as socially desirable (positive) or undesirable (negative) were selected and translated into Japanese. Participants then assessed their deviation from an average peer based on these traits using a standardized visual analog scale ranging from −1 to 1, where −1 indicates feeling inferior to others and +1 indicates feeling superior to others. This approach enabled us to calculate their SI scores. Negative trait ratings for each individual were reversed and combined with their positive trait ratings to determine the average deviation from the midpoint of 0.

### Image acquisition

Structural and functional images were acquired using a 3.0-Tesla Siemens Verio MRI scanner (Siemens Healthcare Sector, Germany) equipped with a 32-channel head coil. Three-dimensional T1-weighted (T1w) structural images were obtained using a magnetization-prepared rapid gradient echo with the following parameters: repetition time (TR) of 2.3 ms; echo time (TE) of 1.95 ms; flip angle (FA) of 9°; slice thickness of 1 mm; field of view (FOV) of 256 × 256 mm^2^; matrix size of 240 × 240; and isotropic voxel size of 1 × 1 × 1 mm^3^. For resting-state functional images, an echo-planar imaging sequence was employed with the following parameters: TR of 2000 ms; TE of 25 ms; FA of 90°; 36 slices; FOV of 220 × 136 × 220 mm^3^; matrix size of 72 × 74; voxel size of 3.0 × 3.0 × 3.0 mm^3^; reconstructed voxel size of 1.72 × 1.72 × 3 mm^3^, and a scan duration of 5 min. During imaging, participants were instructed to close their eyes, relax, refrain from engaging in specific cognitive tasks, and avoid falling asleep.

### Structural magnetic resonance imaging analyses

VBM analysis was conducted using the standard Diffeomorphic Anatomical Registration Through Exponentiated Lie Algebra (DARTEL) (11) processing pipeline in Statistical Parametric Mapping software (SPM12; Wellcome Trust Centre for Neuroimaging, London, UK) with MATLAB 9.10.0 (MathWorks, USA). All images underwent thorough quality control to identify and address artifacts. Each image was spatially normalized to the Montreal Neurological Institute (MNI) coordinate system. T1-weighted scans were segmented into gray matter (GM), white matter (WM), cerebrospinal fluid (CSF), and skull compartments using the segmentation method in SPM12. The DARTEL algorithm was employed to create study-specific templates, and GM images were normalized to the MNI space, resampled to 1.5 mm isotropic voxels, and smoothed using an 8 mm full-width at half maximum (FWHM) Gaussian kernel. Total intracranial volume (TIV) was estimated using the Tissue Volumes utility in SPM12.

We performed multiple regression analyses to identify brain regions associated with SI scores. In these analyses, SI scores served as the independent variable, whereas gender, age, and TIV were included as covariates to control for potential confounding factors. To specifically address variations in SI scores attributable to aging, our analyses utilized age-adjusted residual SI scores as the independent variable. Statistical significance was determined at a voxel-wise threshold of uncorrected P < 0.0001, with an extent threshold of 10 voxels. Significant clusters were used as regions of interest (ROIs) for subsequent resting-state functional connectivity (rsFC) analysis. GM volumes from these ROIs were extracted and plotted against the SI score for visualization.

### Resting-state fMRI analyses

Resting-state fMRI data were analyzed using the CONN toolbox (version 18b; Whitfield-Gabrieli & Nieto-Castanon, 2012, [www.nitrc.org/projects/conn]). Data preprocessing involved several steps. Initially, the first 15 scans were excluded. Realignment and unwarping corrected for subject motion, whereas slice-timing correction addressed interslice differences in acquisition time. The Artifact Detection Tools method identified outlier scans for removal. Segmentation and normalization simultaneously segmented GM, WM, and CSF and normalized the structural data to MNI space. Subsequently, spatial smoothing was performed using a 6mm FWHM Gaussian kernel. Denoising comprised bandpass filtering (0.008–0.09 Hz) to mitigate linear drift and high-frequency noise, alongside confounding factors such as WM and CSF signals, six motion parameters and their derivatives, and scrubbing.

Our statistical analysis framework employed multiple linear regression models to explore associations between SI scores and rsFC within the elderly cohort. Furthermore, differences in seed-to-voxel rsFC associated with SI scores across age groups were assessed. Functional connectivity analyses utilized clusters identified via VBM as ROIs, with the mean time series for each ROI serving as predictors during seed-to-voxel general linear model analysis. Statistical significance was determined using a voxel-wise threshold of p < 0.001 (uncorrected) and a cluster-level correction threshold of p < 0.05 (false discovery rate corrected), with age and gender as covariates (12). For visualization, SI scores were represented as z-scores plotted against the strength of rsFC (Fisher-transformed correlation coefficients) among ROIs, demonstrating significant associations.

## Results

### Behavioral results

SI score analysis revealed the presence of SI across all age groups. The mean SI score was 0.15 ± 0.18, suggesting that our participants exhibited a prevailing sense of superiority regarding their abilities and traits across the age spectrum. Furthermore, a positive correlation was observed between age and the degree of superiority (Pearson’s r = 0.21; p < 0.05), indicating a tendency for SI to intensify with age (Fig. 1).

**Fig. 1.**
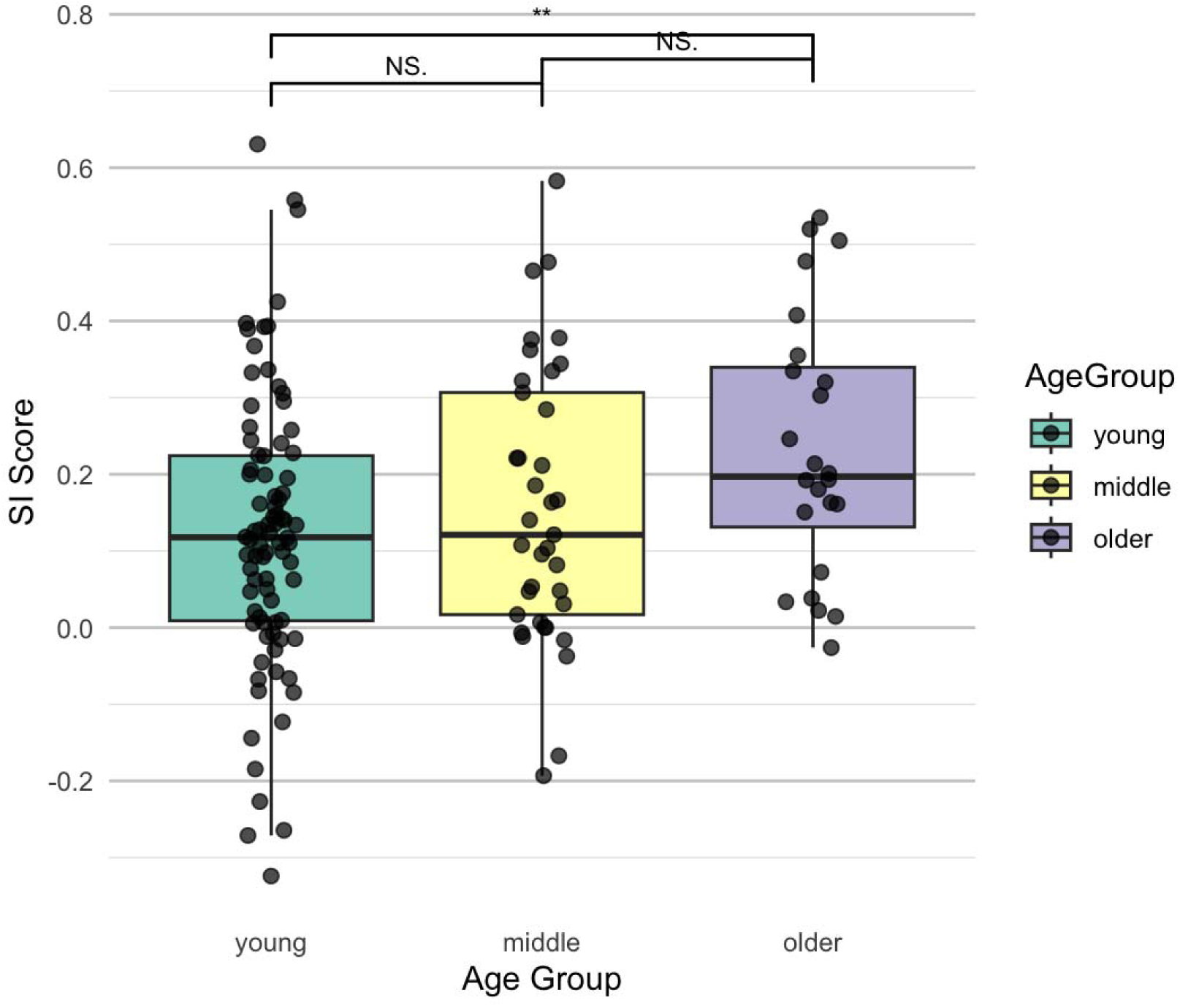
Comparison of the superiority illusion scores across different age groups

### Structural brain regions linked to SI score in older individuals

A comparison between the older group and a combination of the young and middle-aged group found a significant positive correlation between SI score and GM volume in the right precuneus (x = 10.5, y = −76.5, z = 48; T = 3.82; voxel-level p < 0.0001; cluster size = 70; Fig. 2a). The relationship between SI score and right precuneus volume across all age groups is presented in Fig. 2b. However, no significant correlations were found between SI score and GM volume in other brain regions.

**Fig. 2.**
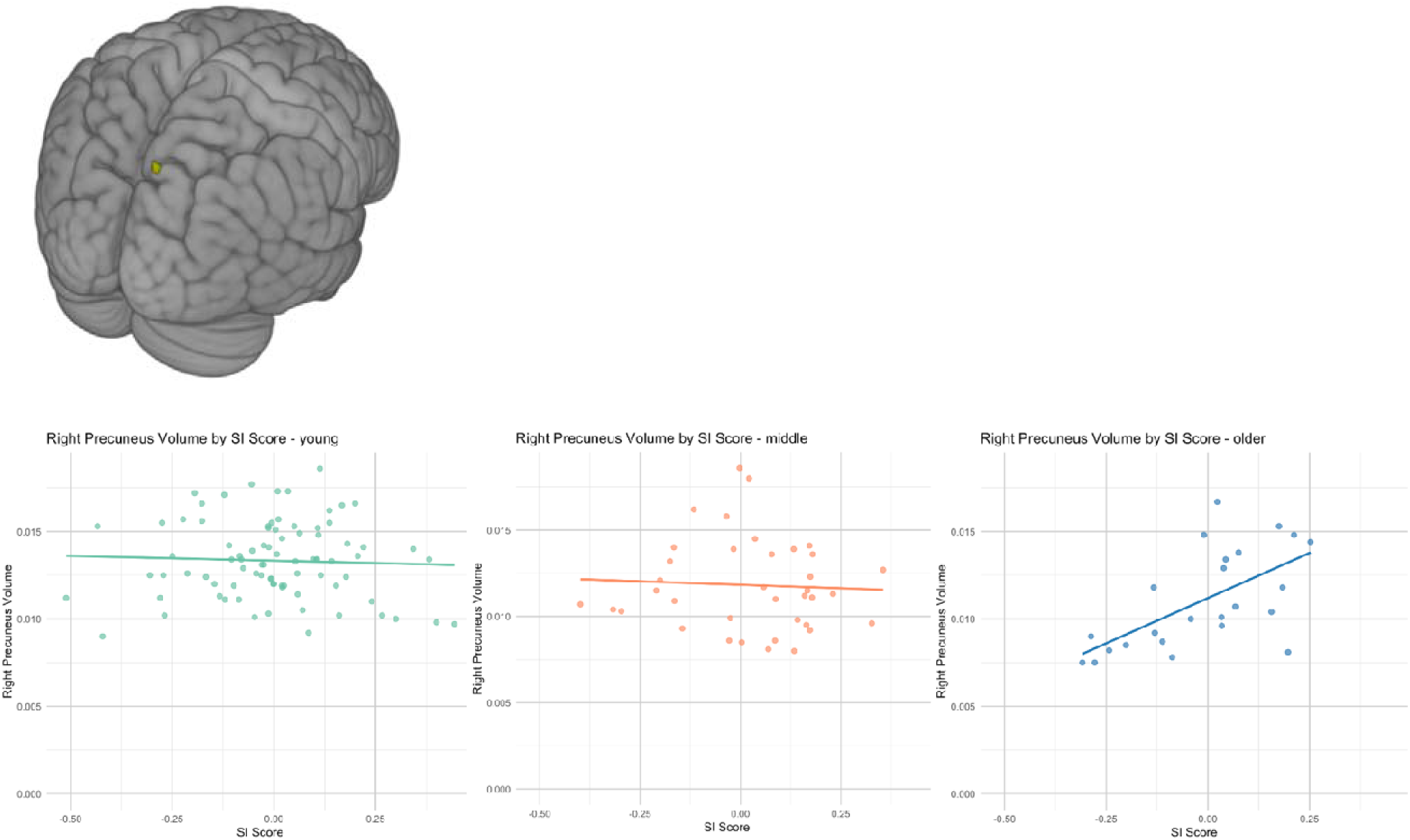
Superiority illusion in healthy older subjects was associated with higher gray matter volume of the right precuneus a) Higher gray matter volume of the right precuneus was associated with superiority illusion b) Scatter plot representation of the superiority illusion effect in the three groups (regression lines are provided for visualization only.)

### rsFC analysis

Seed-to-voxel analysis was employed to investigate the relationship between the SI score and rsFC, focusing on the functional connectivity associated with the right precuneus cluster identified during VBM analysis. Although no associations were observed between SI scores and rsFC in the young- or middle-aged groups, a significant negative correlation was observed between SI scores and connectivity within the left lateral occipital cortex among older participants. These findings have been illustrated in a graph plotting the SI score against the connectivity strength (Fig. 3).

**Fig. 3.**
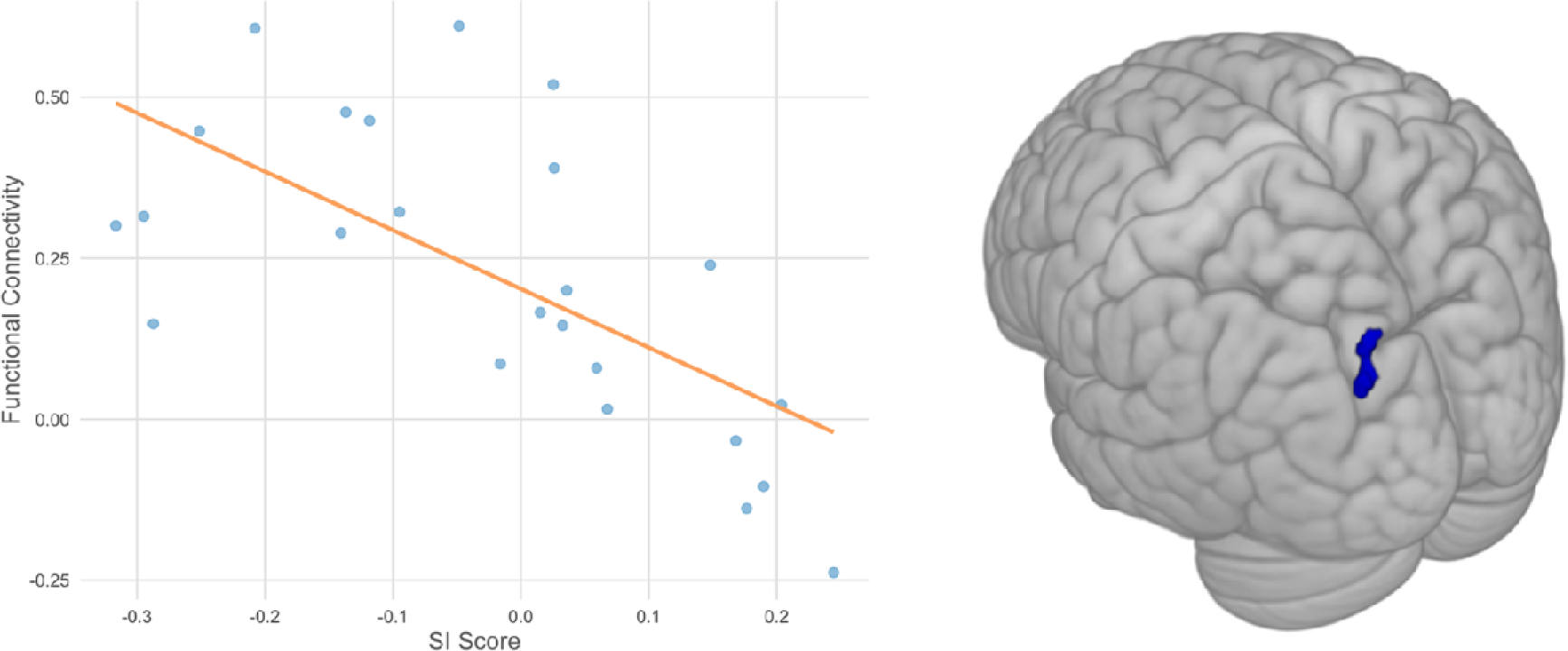
Association between superiority illusion scores and resting-state functional connectivity (rsFC) linking the right precuneus and left occipital cortex in the older group MNI = −24 − 88 + 36 Y axis: rsFC strength between the right precuneus and left lateral occipital cortex (z score) For illustrative purposes, SI scores were represented as z-scores plotted against the strength of rsFC (Fisher-transformed correlation coefficients).

## Discussion

The current study found an increase in the inclination toward the SI with age. Moreover, among healthy older adults, the right precuneus volume correlated with the propensity for the SI. This association was not evident among the middle-aged or younger groups. Furthermore, a weaker rsFC between the right precuneus and visual cortex was observed among elderly individuals exhibiting higher levels of the SI.

Our finding of a more evident SI among older adults can be understood within the framework of the socioemotional selectivity theory, which posits that people prioritize emotionally meaningful goals and experiences as they age, leading to a greater focus on positive information (13). This shift helps older adults maintain a positive self-perception and emotional well-being despite the challenges and losses associated with aging (14, 15). Brain imaging studies have further revealed that older adults show increased frontal lobe activity when processing positive information and increased brain reward system activity in response to positive stimuli (16). These brain response patterns and the tendency to exhibit a positive bias suggest that the SI may serve as a cognitive strategy for maintaining a positive self-perception, thereby contributing to emotional well-being and resilience despite aging-related challenges.

The observed positive correlation between the SI and right precuneus volume may indicate that this region is critical in maintaining positive self-perceptions in older adults. This finding is consistent with the notion that the precuneus plays a role in self-awareness and introspection—processes crucial for maintaining a positive self-view (17). The aforementioned finding also supports the idea that the precuneus is involved in positive affect and life satisfaction, thereby contributing to the experience of positive emotions and contentment with life (18). Moreover, the decreased rsFC between the right precuneus and lateral occipital cortex among older adults with higher SI suggests reduced integration of external visual information during self-referential processing (3, 19). This reduced integration may bias cognition toward more favorable self-perceptions, given that less integration of external feedback could reinforce an individual’s positive self-image.

These findings align with the broader literature on aging and brain function (18, 20–22). The concept of network reorganization in aging suggests that the brain may alter connectivity patterns to compensate for declines in certain cognitive functions (21, 22). This compensation could manifest as increased activation in other brain areas or altered connectivity, which could help maintain cognitive function despite structural declines. In our study, the preserved volume of the right precuneus, despite reduced connectivity, may represent a compensatory mechanism that supports the maintenance of positive biases like the SI.

Although the SI may serve as an adaptive psychological mechanism to preserve well-being, it also highlights the potential vulnerability of older adults to cognitive biases that could influence their decision-making and social interactions. Future research should explore the interplay between structural brain changes, functional connectivity, and cognitive biases in aging populations to develop targeted interventions that enhance mental health and quality of life among older adults.

Although the current study provides valuable insights, several limitations must be considered. Initially, the small sample size of older participants may limit the generalizability of the findings. Hence, future studies with more extensive and diverse samples must confirm our findings and explore potential cultural differences. Our study did not account for several potentially influential factors, such as cognitive functioning, educational background, health status, lifestyle choices, and psychological stress. Thus, future research should consider these variables to provide a more comprehensive understanding of the factors influencing the SI and its neural correlates. Finally, the cross-sectional nature of this study precludes conclusions about causality. Longitudinal studies are therefore necessary to determine the temporal dynamics of these associations.

## Conclusion

Taken together, the current study highlights the critical role of the right precuneus in the positive distortion of self-perception associated with aging. Our findings contribute to a growing body of literature on the neural correlates of positive cognitive biases and their adaptive functions in older adults. Nonetheless, further research is needed to deepen our understanding of these phenomena and develop strategies that enhance the aging population’s mental health and overall well-being.

## Authors’ contributions

Yuki Shidei: Writing – review & editing, Writing – original draft, Project administration, Methodology, Investigation, Formal analysis, Data curation, Conceptualization. Daisuke Matsuyoshi: Writing – review & editing, Project administration, Methodology, Investigation, Formal analysis, Data curation, Conceptualization. Ayako Isato: Writing – review & editing, Project administration, Data curation. Genichi Sugihara: Writing – review & editing, Project administration, Methodology, Investigation, Formal analysis, Conceptualization. Hidehiko Takahashi: Writing – review & editing, Supervision, Project administration, Methodology, Investigation, Formal analysis, Conceptualization. Makiko Yamada: Writing – review & editing, Supervision, Project administration, Methodology, Investigation, Funding acquisition, Formal analysis, Data curation, Conceptualization.

## Acknowledgments

We would like to thank enago (https://www.enago.jp/) for the English-language editing of the manuscript.

## Ethics approval statement

The study was approved by the Committee of Ethics, National Institutes for Quantum Science and Technology, Japan.

## Patient consent statement

The data used in this study were collected under comprehensive consent for future research.

